# From plants to ants: Fungal modification of leaf lipids for nutrition and communication in the leaf-cutter ant fungal garden ecosystem

**DOI:** 10.1101/2020.07.28.224139

**Authors:** Lily Khadempour, Jennifer E. Kyle, Bobbie-Jo M. Webb-Robertson, Carrie D. Nicora, Francesca B. Smith, Richard D. Smith, Mary S. Lipton, Cameron R. Currie, Erin S. Baker, Kristin E. Burnum-Johnson

## Abstract

Lipids are essential to all living organisms, as an energy source, as an important cellular structural component, and as a communication tool. In this study, we used global lipidomic methods to evaluate the lipids in leaf-cutter ant fungal gardens. Leaf-cutter ants and their coevolved fungal cultivar, *Leucoagaricus gongylophorus*, are a model mutualistic system. The fungus enzymatically digests fresh plant material that the ants cut and deliver, converting energy and nutrients from plants, and providing them to the ants through specialized hyphal swellings called gongylidia. Using combined liquid chromatography, ion mobility spectrometry, and tandem mass spectrometry we evaluated differences between the molecular speciation of lipids in the leaf-cutter ant fungal garden ecosystem. This lipidomic study characterized leaves that are fed to the gardens, gongylidia that are produced by the fungus to feed the ants, and spatially resolved regions of the fungal garden through stages of leaf degradation. Lipids containing alpha-linolenic acid (18:3) were enriched in leaves and the top of the gardens, but not dominant in the middle or bottom regions. Gongylidia were dominated by lipids containing linoleic acid (18:2). To evaluate the communicative potential of the lipids in fungal gardens we conducted a behavioral experiment that showed *Atta* leaf-cutter ants responded differently to 18:3 and 18:2 fatty acids, with aggression towards 18:3 and attraction for 18:2. This work demonstrates the role of lipids in both the transfer of energy and as an inter-kingdom communication tool in leaf-cutter ant fungal gardens.

**Importance:** In this work we examined the role of lipids in the mutualism between leaf-cutter ants and fungus. These ants cut fresh leaf material, which they provide to their fungal cultivar, that converts energy and nutrients from the plants and provides it to the ants in specialized hyphal swellings called gongylidia. This work constitutes the first example of a global lipidomics study of a symbiotic system and provides insights as to how the fungus modifies plant lipids into a usable source for the ants. Through a behavioral experiment, this work also demonstrates how lipids can be used as an inter-kingdom communication tool, in this case an attractant, rather than as a repellant, which is more often seen.

## Introduction

Lipids are one of four fundamental components of organisms, the other three being proteins, carbohydrates, and nucleic acids. Lipids serve many roles, and their characterization is critical to understanding how organisms function individually, and in communities. Two roles that we are particularly interested in are as a source of energy and their use in interspecies communication. Lipids are widely used as a form of stored energy in organisms, and in diets represent a dense source of energy, approximately double that of either proteins or carbohydrates (1).

The role of lipids in communication is vital across biological scales. Lipids found in cell membranes are important in cell-to-cell communication, either as a means to identify and differentiate between cells of the same multicellular organism (2), conspecifics in unicellular organisms (3), or cells that do not belong – pathogens (4), predators (5), or food (6). Lipids even direct communication between host cells and virus, thereby impacting the favorability of membrane fusion events (7). At a larger scale, lipids are used as signaling molecules in the form of hormones (between cells of the same organism), pheromones (intraspecific communication), and allelochemicals (interspecific communication).

Previous work on allelochemicals in symbiosis has focused largely in the realm of plant defense against herbivory. When consumed by insect herbivores, some plants emit allelochemicals that attract the herbivore’s natural enemy, parasitic wasps (8). This system is a fascinating example of communication in a multipartite symbiosis between a host, herbivore, and indirect defensive symbiont. We expect that this type of communication would exist in beneficial symbioses as well. Our work here explores the role of lipids, in both nutrition and communication, in a model mutualistic system where fungi serve as an interface between ants and plants.

Leaf-cutter ants are dominant herbivores that can consume as much as 17% of the leaf biomass produced in Neotropical ecosystems (9). *Atta* leaf-cutter ant colonies contain millions of workers that fall into specialized casts (10), contributing to the colony’s success as ecosystem engineers (11). Foraging leaf-cutter ants are often seen marching in conspicuous trails and bringing pieces of leaves back to their underground nests that can be as large as 30 m^2^ to 35 m^2^ in area and several meters deep. Once to the nest, the ants use the leaves to feed their fungal cultivar, *Leucoagaricus gongylophorus* (12), which is grown in their specialized fungal gardens. By cultivating and feeding these fungal gardens, the ants are able to access nutrients in plant biomass that would otherwise be unavailable (13–17).

Leaf-cutter ants and their microbial symbionts are paradigmatic in the field of insect microbial symbiosis. Unlike other herbivores, the microbes in symbiosis with leaf-cutter ants can be easily sampled because they exist as an external digestive system in the form of fungal gardens. These fungal gardens exhibit vertical stratification, characterized by top, middle and bottom regions. From top to bottom, there is a visual gradient of leaf degradation with leaves starting as fresh, undigested leaves at the top, then processing through the initial stages of degradation in the middle, and finally to more complete stages of degradation by the bottom (18). To promote growth of the fungal garden, leaf-cutter ants perform different activities in the various locations of the garden. Specifically, the ants triturate fresh plant material, incorporate it into the top of the fungal garden, deposit tufts of fungal mycelium, and defecate with excrement containing fungus-derived plant biomass-degrading enzymes to initiate lignocellulose degradation (19, 20). As the plant material placed on the top of the fungal garden is digested, it moves toward the bottom of the garden where it is removed from the fungal garden chamber and deposited in a refuse dump by the ants (21), and by which time it has been depleted of most nutrients. The gradual degradation of the plant material from top to bottom of the garden, combined with specific ant activities, gives each region of the fungal garden a different visual appearance and varied molecular properties (**Figure 1**). The middle of the fungal garden, where the density of fungal hyphal growth is most pronounced, is also where the *L. gongylophorus* produces gongylidia, or specialized hyphal swellings, that worker ants consume and feed to larvae in the colony. While foraging workers obtain some nutrition from the plant sap they drink when they cut the fresh leaf material, garden workers and larvae exclusively consume gongylidia, where they obtain carbohydrates, polysaccharides, and energy-rich lipids (22, 23).

**Figure 1:**
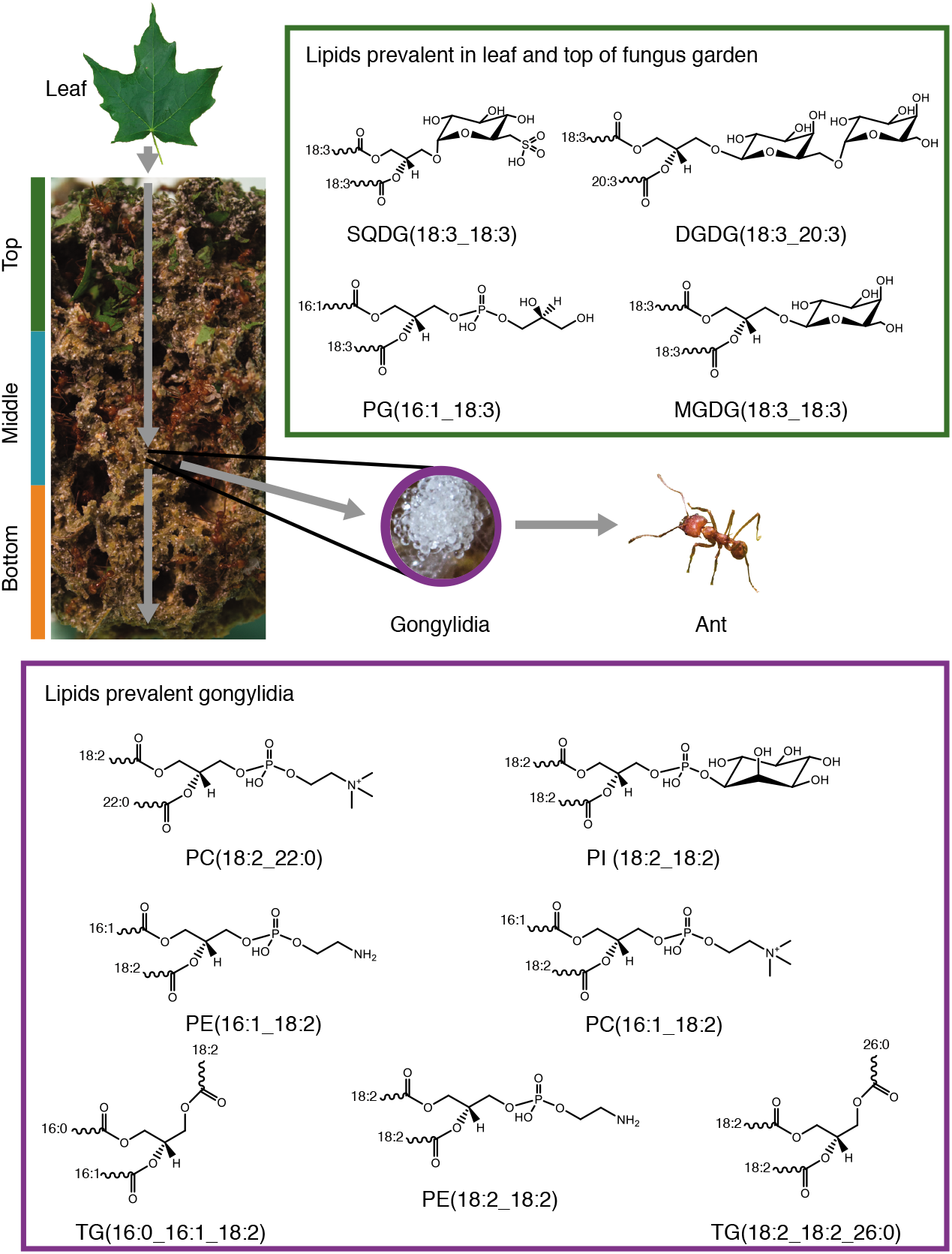
A schematic diagram of the process by which lipids are processed through the fungal garden. Fresh leaf material is first deposited on the top of the fungal garden and passes through the middle and bottom of the garden. As the leaf material proceeds, lipids are extracted by the fungus and modified. The final lipid profile in the gongylidia (the specialized fungal structure that ants consume) differs from that which is in the leaf material and top of the fungal garden. The gongylidia are characterized by lipids containing linoleic acid (18:2), whereas the leaves are characterized by lipids containing alpha-linolenic acid (18:3). Lipid structures here are only representative and other isomers are possible. Photo credits: Lily Khadempour, except gongylidia by Don Parsons.

To elucidate the relationship between leaf-cutter ants and their fungal cultivar, earlier studies have focused on understanding how the fungus breaks down plant biomass (13, 14). In our previous metaproteomic analyses of the top, middle and bottom regions of the fungal garden, we discovered that *L. gongylophorus* produced a majority of the lignocellulases found (13). The studies also showed the resident bacterial community likely aids in this process by producing amino acids and vitamins, thereby enabling the fungus to thrive (13, 14, 24). These results shed great light into the symbiosis between the ant and the fungal cultivar, but to understand a more complete picture other proteins were evaluated. Interestingly, the gongylidia (13, 20) showed an enrichment in unique lipid-associated proteins when compared to different regions of the garden (Data file S1), ultimately suggesting the gongylidia lipids would differ from those throughout the fungal garden. A detailed analysis of the molecular speciation of lipids is needed to provide biological insight into subpopulations and differential activities of complex samples, such as the fungal garden ecosystem.

Here, we conduct the first global lipidomic study of the leaf-cutter ant-microbe symbiosis using comprehensive liquid chromatography–tandem mass spectrometry and liquid chromatography–ion mobility spectrometry-mass spectrometry. We examined spatiotemporal changes in the molecular speciation of lipids across six heterogenous *Atta* leaf-cutter fungal gardens through different stages of leaf degradation. First we evaluated the lipid content of leaves the ants use to feed their cultivar to understand which ones existed initially. Then we assessed the gongylidia, and the top, middle, and bottom regions of their fungal gardens at initial, intermediate, and advanced stages of leaf degradation, respectively (**Figure 1**). The lipid content of the leaf material was compared to the different regions of the fungal garden to understand consumption of the leaf lipids and synthesis of the fungal garden consortium lipids through the various regions. Additionally, the lipid content of the gongylidia was compared to the middle region of the fungal garden to evaluate its specific properties versus the area where it was harvested. Upon finding an enrichment of particular lipids in the leaves and fungal garden components, we conducted a behavior experiment using seven *Atta cephalotes* colonies to observe whether worker ants detected and responded to the lipids differently.

## Results

We observed that the predominant lipids in the gongylidia and top and bottom of the garden varied greatly. The top of the garden was most similar to the leaves and had predominant lipid subclasses including monogalactosyldiacylglycerol (MGDG), sulfoquinovosyldiacylglycerol (SQDG), and diacylglycerophosphoglycerol (PG), while the lipids in the bottom of the gardens included diacylglycerophosphoethanolamines (PE) and diacylglycerophosphoserine (PS). The gongylidia had dominant lipids including ceramides (Cer), diacylglycerophosphoethanolamines (PE), and triacylglycerols (TG). Across all lipid subclasses interesting trends were observed based on individual fatty acid composition; an evaluation of the lipid subclasses and fatty acid composition differences is given below.

### Lipid comparisons of leaves and fungal garden regions

To initiate the study, lipids present in the maple leaves fed to the fungal garden were evaluated so they could be compared to the fungal garden regions and gongylidia. The lipids present in the leaves were evaluated and leaf lipid categories included sphingolipids, glycerophospholipids, and glycerolipids. These identifications were dominated by the PG subclass as well as galactolipids (MGDG and SQDG) with only minor amounts of the other phospholipids (diacylglycerophosphocholines (PC), PE, and diacylglycerophosphoinositols (PI)) (see Data file S2 for all characterized lipids). Evaluation of the fungal garden samples showed that 274 lipids were identified from the top, middle, and bottom regions of the six gardens (Data file S2 (a) & (b)). Both the leaf lipids and garden lipids are depicted in **Figure 2** where the relative log_2_ expression level of each lipid is represented by a red-white-blue color scale with red denoting high abundance and blue low abundance. Leaf lipids are included in the top row (highlighted with a pink box), subsequent rows contain garden lipids, for the six top regions (highlighted with a green bar), six middle regions (highlighted with a teal bar), and six bottom regions (highlighted with an orange bar). In leaf samples, PGs and galactolipids were observed to be the most abundant, which was distinct from the lipids detected in the top, middle, and bottom regions of the garden, where TGs were found to be most abundant.

**Figure 2:**
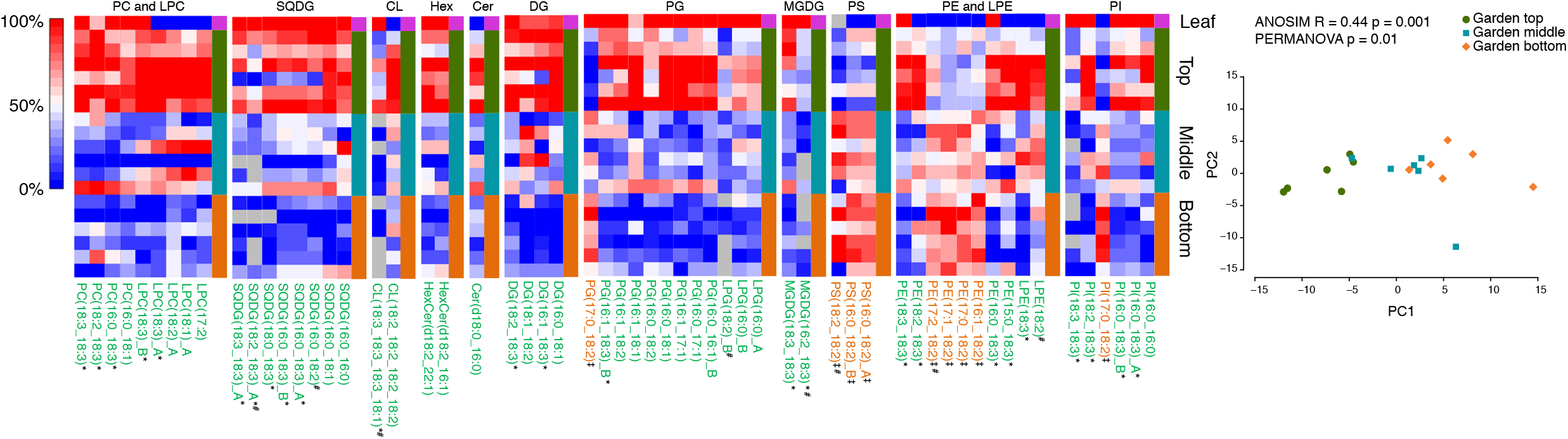
Heatmaps representing the relative log_2_ expression level for lipids in the leaf food source and the top, middle, and bottom strata of six leaf-cutter ant fungal gardens. Each column represents the relative abundance of a unique lipid and the heatmaps are scaled by column. Lipids significantly increased in the top are denoted with green font; lipids significantly increasing in the bottom are denoted with orange font (p-values < 0.05 were deemed significant); These lipids were also significantly different with a Benjamini-Hochberg adjusted p-value threshold of 0.05 with the exception of the eight lipids denoted with #; _A and _B denote structural isomers. Diacylglycerophosphocholines (PC), monoacylglycerophosphocholines (LPC), sulfoquinovosyldiacylglycerol (SQDG), Monogalactosyldiacylglycerol (MGDG), Cardiolipin (CL), phosphatidic acid (PA), Hexosyl-Ceramide (HexCer), Ceramide (Cer), Diacylglycerols (DG), diacylglycerophosphoglycerol (PG), monoacylglycerophosphoglycerol (LPG), diacylglycerophosphoserine (PS), diacylglycerophosphoethanolamines (PE), monoacylglycerophosphoethanolamines (LPE), diacylglycerophosphoinositols (PI). Lipid abbreviations show the total number of acyl chain carbons: total number of double bonds. ‡, 18:2 containing lipids significantly increased in the bottom of the garden; *, 18:3 containing lipids significantly increased in the top of the garden.

Garden lipids were also from sphingolipids, glycerophospholipids, and glycerolipids categories and 13 subclasses. The glycerophospholipid and glycerolipid categories represented 97% of the identified lipids with 89 glycerophospholipids species and 177 glycerolipids species, while the sphingolipid category was only a minor component with 8 species identified in the garden (Data file S2(b)). Figure S1 and Data file S2(c) depict 110 lipids, including isomeric species (25), with accurate quantitative values and no co-eluting lipid species detected. When the top and bottom fungal garden regions were statistically compared to evaluate lipid changes and leaf degradation, lipid species were found to be statistically significant and a trend was noted pertaining to chain length and degree of unsaturation. PE, PG, diacylglycerol (DG), and TG species, were more unsaturated (i.e., contained more double bonds in the fatty acid chains) in the top region of the garden, and PG and TG species had longer fatty acid chains.

An examination of the individual fatty acid composition of the lipids in the fungal garden also illustrated interesting trends. The fatty acyl 16:0, 18:2, and 18:3 groups were found to dominate the fatty acids profiles for the most abundant lipids identified in the phospholipid and glycerolipid subclasses. **Figure 2** details the acyl chains for the 59 significantly enriched lipids (p-value < 0.05; 51 lipids, adjusted p-value < 0.05), where those highlighted in green text (50 lipids) or orange text (9 lipids) are significantly enriched in the top or bottom garden layer, respectively. For example, PC(18:3_18:3), LPC(18:1)_A, PE(18:3_18:3), and PI(18:3_18:3) are highlighted with green text in **Figure 2** since they greatly decreased from the top to bottom region of the fungal garden. Of the 50 lipid species significantly elevated in the top layer, over half of them contained an 18:3 acyl chain. These 18:3 containing lipids significantly more abundant in the top layer include MGDG, SQDG, monoacylglycerophosphocholine (LPC), PC, PG, DG, Cardiolipin (CL), PI, monoacylglycerophosphoethanolamines (LPE), and PE species (signified with an * in **Figure 2**) and TG(18:3/18:3/18:3) (Data file S2(b) denotes the 12 significant (adjusted p-value < 0.05) TGs, where no co-eluting species were detected, increasing in the top layer). Many of these lipids also had a relatively high abundance in the leaves, especially those containing two 18:3 fatty acyl groups (e.g., 18:3_18:3), suggesting 18:3 containing lipids come from the leaves. Only nine lipids were found to significantly increase in the bottom layer of the garden compared to the top and, all of these lipids contained at least one 18:2 (signified with a ^‡^ in **Figure 2**), suggesting 18:3 is degraded from the top to bottom regions of the garden by fungal garden microbial symbionts, while 18:2 is synthesized by the fungal garden. Based on these results we hypothesized that 18:2 could be enriched in the gongylidia.

### Lipid changes between fungal garden and gongylidia

Next, the gongylidia were compared to the middle fungal garden where they are located to assess similarities and differences. In the gongylidia, 263 lipids species were identified from 18 different subclasses (Data file S3 (a) & (b)). Figure S2 depicts 183 lipids, including isomeric species, with accurate quantitative values and no co-eluting lipid species detected (Data file S3(c)). From the statistical comparison of the gongylidia and surrounding middle garden region, **Figure 3** depicts the 116 lipid species found to be statistically significantly enriched (p-value < 0.05; 109 lipids, adjusted p-value < 0.05), indicating the gongylidia lipids are distinct from the surrounding middle region. Both the middle region garden lipids and gongylidia lipids are depicted in **Figure 3** where the relative log_2_ abundance of each lipid is represented by a red-white-blue color scale, red denotes high abundance and blue denotes low abundance. Leaf lipids are included in the top row (highlighted with a pink box), subsequent rows contain garden lipids, for the six middle regions (highlighted with a teal bar), and six gongylidia samples (highlighted with a purple bar). The acyl chains for each lipid are also detailed in **Figure 3**, where those highlighted in purple text or teal text are significantly increasing in the gongylidia or surrounding middle garden region, respectively. Lipid categories with high abundance in leaves from the initial garden study, MDGD, SQDG, and PG, in addition to DGDG, showed low enrichment in the gongylidia samples; whereas, both detected mannosylinositol phosphorylceramide (MIPC) lipids, MIPC(t18:0_24:0(2OH)) and MIPC(t18:0_26:0(2OH)), and all five significant Cer species were enriched in the gongylidia. PC, PE, phosphatidic acid (PA), PI, DG, and TG lipids showed differential enrichment dependent on the lipid fatty acyl composition. All significantly changing 18:3 containing PC, PE, PA, and DG species in the middle garden were depleted in the gongylidia (signified with an * in **Figure 3**). Conversely, all significantly changing 18:2 containing PC, PE, PA, DG, and PI species were enriched in the gongylidia (signified with a ^‡^ in **Figure 3**). Many TG species contained both 18:3 and 18:2 as one of their three fatty acyl chains, yet ninety percent of all TG species depleted in the gongylidia compared to the surrounding fungal garden contain at least one 18:3 acyl chain and >84% of all TG species enriched in the gongylidia contain at least one 18:2 acyl chain. Additionally, lipid subclasses with high abundance in the leaves (MDGD, SQDG, PG, and DGDG) showed relatively low concentrations in the gongylidia samples.

**Figure 3:**
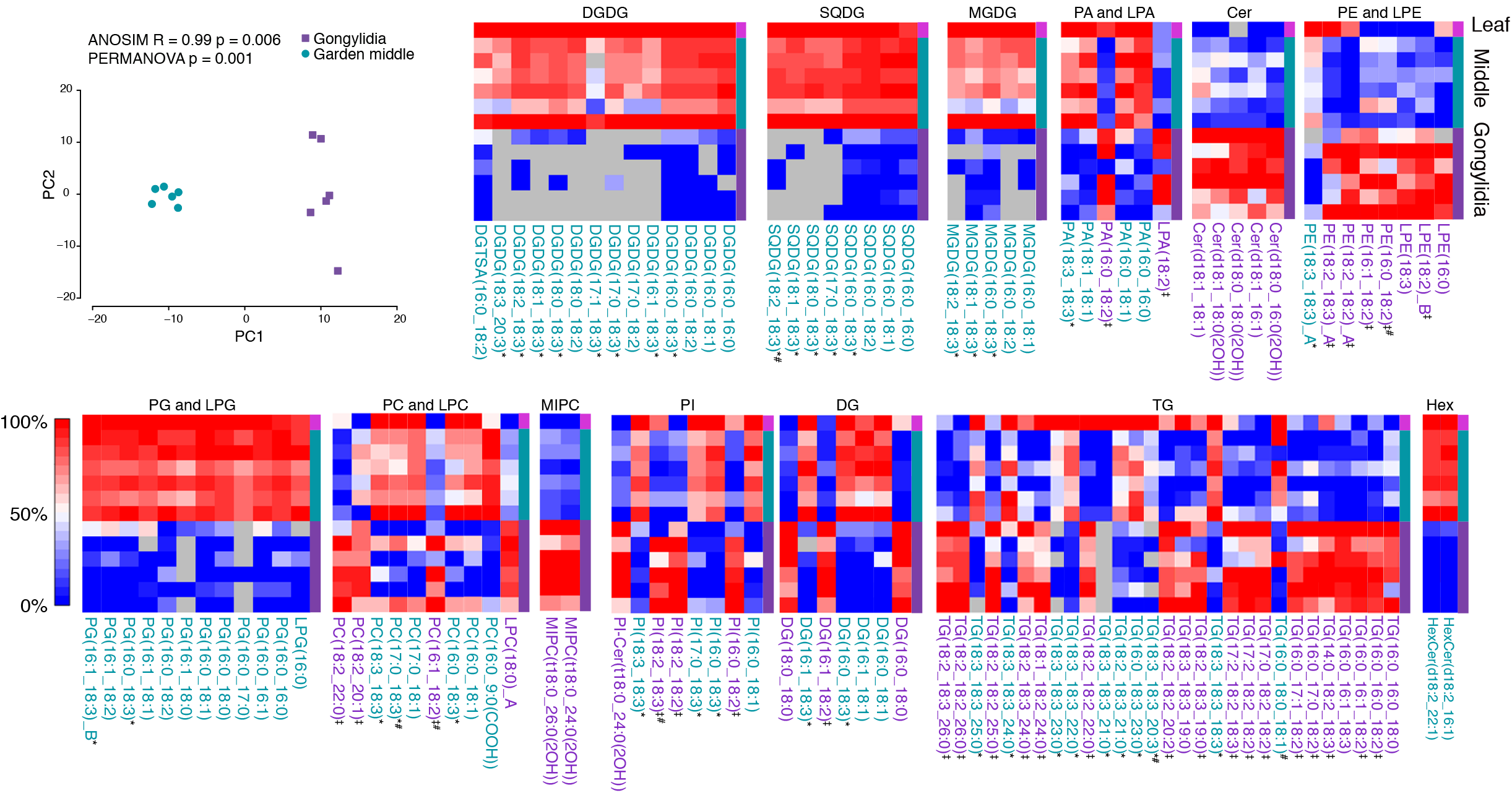
Heatmaps representing the relative log_2_ expression level for lipids in the leaf food source and the gongylidia and surrounding fungal garden of six leaf-cutter ant fungal garden ecosystems. Each column represents the relative abundance of a unique lipid and the heatmaps are scaled by column. Lipids significantly enriched in the gongylidia compared to the surrounding fungal garden are denoted with purple font; lipids significantly enriched in the fungal garden are denoted with teal font (p-values < 0.05 were deemed significant); These lipids were also significantly different at a Benjamini-Hochberg adjusted p-value threshold of 0.05 with the exception of the seven lipids denoted with #;_A and _B denote structural isomers. Digalactosyldiacylglycerol (DGDG), sulfoquinovosyldiacylglycerol (SQDG), Monogalactosyldiacylglycerol (MGDG), diacylglycerophosphoglycerol (PG), monoacylglycerophosphoglycerol (LPG), Diacylglycerophosphocholines (PC), monoacylglycerophosphocholines (LPC), mannosylinositol phosphorylceramide (MIPC), Ceramide phosphoinositol (PI-Cer), diacylglycerophosphoinositols (PI), phosphatidic acid (PA), lysophosphatidic acid (LPA), Hexosyl-Ceramide (HexCer), Ceramide (Cer), diacylglycerophosphoethanolamines (PE), monoacylglycerophosphoethanolamines (LPE), Triacylglycerols (TG), Diacylglycerols (DG). Lipid abbreviations show the total number of acyl chain carbons: total number of double bonds. *, 18:3 containing lipids significantly decreased in the gongylidia; ‡, 18:2 containing lipids significantly increased in the gongylidia.

### Behavior experiment

Our lipidomic analyses characterized a prevalence of lipid species containing linoleic acid (18:2) in the gongylidia, the leaf-cutter ant’s food source, and lipid species containing alpha-linolenic acid (18:3) in leaf material. We performed a behavior experiment to test if the *Atta cephalotes* workers had different responses to untreated paper discs and paper discs treated with oleic acid (18:1), 18:2, and 18:3 (**Figure 4**; Figure S3, S4, & S5; Movie S1 & S2). Overall, the ants spent significantly more time inspecting 18:2 (p=0.0012) and 18:1 (p=0.015) treated paper discs than the control paper discs. Similarly, the ants picked up the discs infused with 18:2 (p=0.039) more frequently than the control discs, but during the experiment never picked up a disc containing 18:3. However, the ants displayed aggressive behavior, including lunging and biting the discs more frequently toward 18:3 compared to the control (p=0.014). The ants did not exhibit aggressive behavior towards the discs with 18:1 or 18:2.

**Figure 4:**
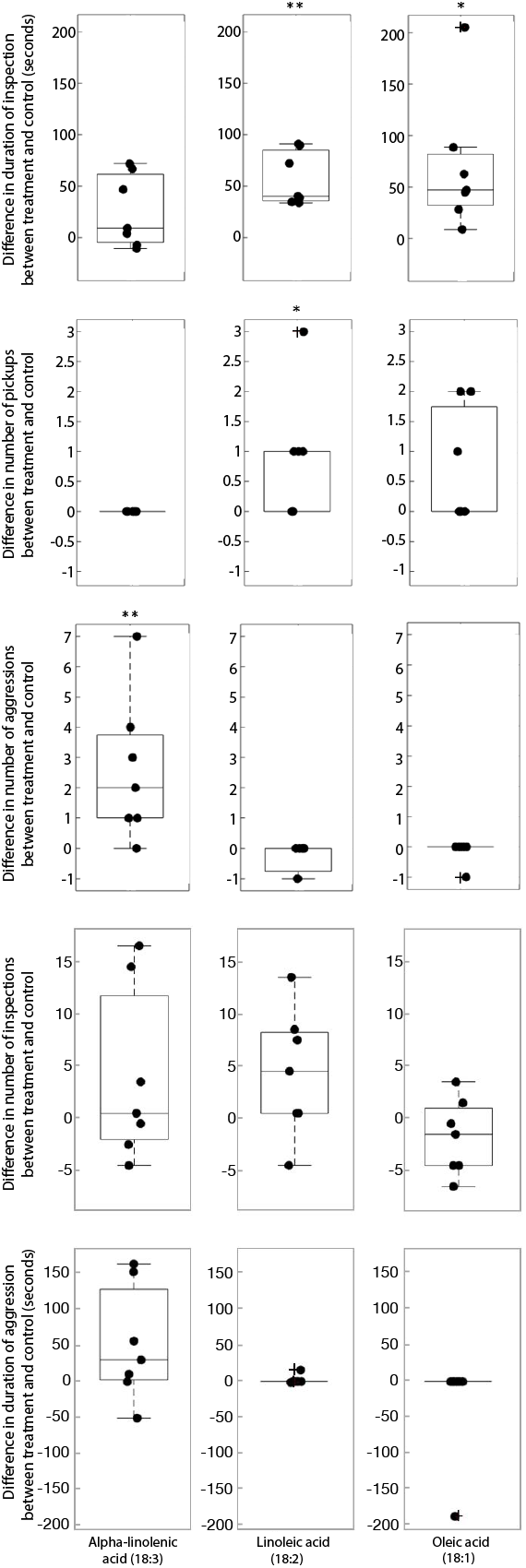
Results of the behavior experiment. *Atta cephalotes* ants are attracted to linoleic acid (18:2) and oleic acid (18:1) but show aggression toward alpha-linolenic acid (18:3). Each point represents one experimental replicate, where the values for the control count or duration are subtracted from the treatment count or duration. A one sample t-test was conducted for the difference in duration and a one sample test of proportion was conducted for the number of pickups and aggressions. Those whose mean is significantly different from zero are marked with * p < 0.05 and ** p < 0.01. Box plot definitions: center line – median, upper and lower box limits – upper and lower quartiles, whiskers – 1.5 x inter-quartile range, outliers – any points outside the 1.5x inter-quartile range.

## Discussion

Leaf-cutter ants, from the genera *Atta* and *Acromyrmex*, cultivate a fungus on fresh foliar material, which converts plant biomass into new forms that are readily usable by the ants and their larvae. This obligate mutualism is derived from within the ant subtribe Attina, which represents a monophylogenetic lineage of ants that have associated with fungal cultivars for approximately 55 million years. The leaf-cutter ants are the most derived group of the Attina and diverged approximately 12 million years ago, coinciding with their association with *L. gongylophorus* (26). Both organisms have undergone co-evolution. *Leucoagaricus gongylophorus* has adapted to its association with the ants and has undergone gene reduction so that it can no longer function as a saprotroph, like its closest free-living relatives (27). On the other hand, the ant’s digestive system has adapted to prevent digestion of fungus-produced plant biomass degradation enzymes so that the ants can redistribute them through fecal fluid onto freshly incorporated leaf material (19). In return, the fungus provides all of the nutrients to feed the colony by producing carbohydrate, polysaccharide, and lipid rich gongylidia (22, 23).

In this study, characterization of both the lipid content of leaf material and lipids synthesized by the fungus, allowed us to gain novel insight into the conversion of lipid nutrients from the leaf material deposited on the top of the fungal garden to the enrichment of lipid nutrients in the gongylidia for the ant to consume. We observed significant subclass trends correlated to each region. There is an enrichment of lipids containing 18:3 in the leaves and the top of the garden, and an enrichment of lipids containing 18:2 at the bottom of the garden and gongylidia. Lipid subclasses that were significantly increased in the top of the garden and leaf material included PG, MGDG, SQDG, and DGDG species, all of which are known to be significant contributors to photosynthetic organelles (28). In addition, 18:3 has notable signaling properties in plant tissues. When leaves are wounded, the polypeptide systemin is emitted from the damaged cells into the apoplast, signaling the liberation of 18:3 from plant membrane lipids into the cells. 18:3 begins the defense pathway by being converted to 12-oxophytodienoic acid and then through beta-oxidation is converted into jasmonic acid (29). The main defense mechanism in these leaves is jasmonic acid, and 18:3 is crucial to its synthesis.

The abundant lipids in the top of the fungal garden through to the gongylidia differ greatly. This is an important result because while the ants ingest biomass-degrading enzymes from the gongylidia and deposit them on top, the lipids are not being recycled in this manner. It is clear then that the fungal garden metabolizes the lipids from the leaves and synthesizes new lipids which are provided to the ants and larvae through the gongylidia, where they are ingested and absorbed into their bodies. While this was assumed previously (22, 23), our study is the first to actually track the molecular speciation and dynamic changes of these lipids through the fungal garden.

We conducted the behavior experiment because we wanted to evaluate whether the ants were able to detect the various lipids in the fungal gardens, and to observe any respective differences in their behavior. We expected *Atta cephalotes* workers to be attracted to all three tested lipids. Previous studies had shown that ants from the genera *Pogonomyrmex, Solenopsis*,and *Atta* are attracted to 18:1, which is a lipid that dead ants emit from their bodies as a cue for workers to move them out of the colony or toward the dump (30–32). We observed similar attractive results with our ants, validating our methods. We expected that the workers would also be attracted to 18:3, which is abundant in leaves they routinely cut and incorporate into their gardens. However, the workers instead behaved aggressively toward 18:3 by lunging and biting at the infused discs indicating that it is not this component of leaves that leaf-cutter ants are attracted to. This ant response to the 18:3 lipids might be related to the lipid’s role as a constituent of the plant defense response lipid, jasmonic acid, which may be playing a role in deterring herbivory by leaf-cutter ants. Finally, we also expected that the *Atta* workers would be attracted to 18:2 since it has been shown attractive to another ant species (*Cataglyphis fortis*) (33) and is abundant in the gongylidia they consume. The ants’ attractive responses to 18:2 (both disc pickup and time spent inspecting) differed most greatly from the controls, compared to the two other lipids tested, which suggests that there is a strong attraction to this lipid. These responses all relate back to the fungal garden metabolizing 18:3 containing lipids from leaves to synthesize 18:2. 18:3 is repulsive to the ants, while 18:2 is highly attractive to them. One very clear example of coevolution between the leaf-cutter ants and *L. gongylophrous* is the cultivar’s morphological and physiological adaptation in the production of gongylidia for ant consumption (34). Our result, indicating that the gongylidia contain lipids that the leaf-cutter ants are attracted to, provides a further example of the intricacies of coevolution through lipid communication in this system. By restricting the enrichment of 18:2 lipids to the gongylidia, the fungus can focus the ants’ consumption to these specialized structures, thereby preventing damage to its hyphae. Such communication from the fungus to the ants is vital for maintaining fungal fitness, and for maintaining this mutualism.

The study of fungal allelochemicals that influence insect behavior is not new, but we have taken a different approach in addressing this topic. Typically, behavior is observed between an insect and a fungus, and then work is done to identify the allelochemicals involved in that communication (35). Often, after identifying these allelochemicals, they are used as tools in pest management (36). In the leaf-cutter ant literature as well, the approach has been largely from the direction of behavior to mechanism, and it has been suggested that the cultivar uses volatile semiochemicals to communicate with the ant colony (37) but attempts at identifying these have not been conclusive (38, 39). Our approach was different. We were interested in identifying and quantifying lipids in the fungal garden, and then based on proteomic and lipidomic data, hypothesized that these lipids could be used as allelochemicals. Our behavioral experiment then supported this and provides a new tract of study for understanding communication between leaf-cutter ants and their cultivar.

As an important constitute of food, lipids act as a valuable source of energy, cellular structural component, and as a communication tool. Thus, understanding what lipids are consumed by leaf-cutter ants and how their cultivar functions to obtain, modify and deliver plant lipids, further sheds light on this complex mutualistic relationship between these organisms. In a broader context, it provides an important example of how microbes mediate the relationship between herbivores and the plants that they consume, either directly, with microbes interfacing with their hosts in the gut, or less directly, with the fungal garden serving as the ants’ external gut. Future lipidomic studies may help us to further understand how these lipids are used by the ants in communication and symbiont maintenance, if and how they are incorporated into the ants’ bodies, and if specific lipids play a role as allelochemicals throughout the Attina and other fungus farming insects, or if this phenomenon is limited to the leaf-cutter ants and their cultivar *L. gongylophorus*.

## Materials and Methods

### Material collection

Six lab-maintained *Atta* leaf-cutter ant colonies were provided exclusively maple leaves for the two weeks before sample collection. We collected leaf material as well as fungal garden from the top, middle, and bottom of leaf-cutter ant fungal garden ecosystems. These layers are differentiated based on color, texture and location in the garden (**Figure 1**). All of the six fungus gardens that were sampled were maintained in the same size of container, 8 x 8 x 8 cm and so the fungus garden that was sampled was from approximately the same distance from the edge for each sample type.

Gongylidia are difficult to collect because of their small size and their sparseness in the garden. To collect gongylidia, we first collected fungal garden in deep petri dishes and reduced the number of workers. With fewer workers in the fungal garden, the gongylidia proliferate. After several days, the gongylidia were picked off by needlepoint under a dissecting microscope and they were placed in a microcentrifuge tube with water.

The leaf material used in this experiment was collected in the summer and kept frozen at −20°C in vacuum sealed bags so that the ant colonies can be fed during the winter when fresh leaves are not available.

### Total lipid extraction

The collected material was lyophilized for untargeted lipidomics analysis. Lipids were extracted using a modified Folch extraction (40). To break up the biological material, approximately 25 mg of sample was bead beaten using a 3 mm tungsten carbide bead in 750 μl of methanol for 2 min at a frequency of 30 Hz. The sample was then removed and transferred into a 20 mL clean EPS glass vial with a Teflon lined cap. Another 750 μl of methanol was added for a final volume of 1.5 mL methanol. Next, 3 mL of chloroform and 200 μl of water were added to each sample. The samples were vortexed for 30 s, sonicated for 30 min, vortexed again for 30 s, and then 0.925 μl of water was added to induce a phase separation. The samples were incubated at 4°C overnight and then the lower lipid layer was removed, dried down, and stored at −20°C at a concentration of 2 μg/μl in 2:1 chloroform/methanol until Mass Spectrometry (MS) analysis.

### Liquid chromatography (LC)-tandem mass spectrometry (MS/MS) and LC-ion mobility spectrometry-mass spectrometry (LC-IMS-MS) analyses

All extracted lipids in this manuscript were analyzed by LC-MS/MS using a Waters NanoAquity UPLC system interfaced with a Velos Orbitrap mass spectrometer (Thermo Scientific, San Jose, CA) and an Agilent 6560 Ion Mobility QTOF MS system (Agilent, Santa Clara, CA). The total lipid extracts (TLE) were reconstituted in methanol for a final abundance of 0.4 μg TLE/μl. Seven μl of the TLE were injected onto Waters column (HSS T3 1.0 mm x 150 mm x 1.8 μm particle size). Lipids were separated over a 90 min gradient elution (mobile phase A: ACN/H_2_O (40:60) containing 10 mM ammonium acetate; mobile phase B: ACN/IPA (10:90) containing 10 mM ammonium acetate) at a flow rate of 30 μl/min. Samples were analyzed in both positive and negative ionization using HCD (higher-energy collision dissociation) and CID (collision-induced dissociation) to obtain high coverage of the lipidome. Velos Orbitrap MS abundance data is depicted in **Figure 2**, **Figure 3**, Data file S2, and Data file S3. Figures and files apply LIPID MAPS Classification, Nomenclature and Shorthand Notation for MS-derived Lipid Structures (41).

The LC-IMS-MS analyses were also performed in both positive and negative ion mode and collected from 100-3200 m/z at a MS resolution of 40,000. The LC-IMS-MS data were analyzed using in-house PNNL software for deisotoping and feature finding the multidimensional LC, IMS and MS data (42).

### Lipid targeted database and alignment

We used LIQUID software (43) for lipid targeted database alignment. This is achieved by aligning all datasets (grouped by sample type and ionization mode) and matching unidentified features to their identified counterparts using MZmine2 (44). Aligned features are manually verified and peak apex intensity values are exported for statistical analysis.

### Behavior experiment

An ant behavior experiment was conducted in order to determine if *A. cephalotes* ants could detect various lipids found in the fungal garden, and to observe their corresponding behavioral response to them. We tested the ants’ response to linoleic acid (abundant in the gongylidia, 18:2 fatty acids), alpha-linolenic acid (abundant in the leaves and top of garden, 18:3 fatty acids), and oleic acid (excreted by dead ants, 18:1 fatty acids). To test the response of the *A. cephalotes* to these lipids, we used a method similar to those from López-Riquelme *et al*. (32). All three of the lipids were dissolved in ethanol at 1 mg/mL concentration. We pipetted 5 μL of the mixture onto 6 mm paper discs (Whatman 2017-006), that had been pre-labeled with pencil, then waited several seconds for the ethanol to evaporate before introducing the discs to the *A. cephalotes* colonies. Discs with only ethanol were used as a negative control. Each treatment replicate was set up as a binary choice between the treatment oil, and the negative control. We used seven *A. cephalotes* queenright colonies, which were all collected from La Selva Biological Research Station in Costa Rica in April 2018 and were maintained at the University of Wisconsin-Madison. The colonies were housed in 42.5 cm x 30.2 cm x 17.8 cm plastic containers. Three treatment discs and three control discs were used for each assay and were placed on a flat surface in the colony box. The ants were observed for five minutes from the time the discs were placed in the colony. Behavior was coded using the BORIS program (45). We recorded the number of times discs were picked up, characterized by ants using their mandibles to lift the disc in the same manner that they lift leaf pieces when foraging. We recorded the number and duration of ant inspections, characterized by ants approaching the discs while waving antennae toward the discs, touching the disks with their antennae, or walking over the discs. Finally, we recorded the number and duration of aggressive behaviors toward the discs, characterized by lunging and biting, or moving slowly or stopping nearby with open mandibles (46) (example behaviors can be seen in Movie S1 and Movie S2). It should be noted that if several ants were simultaneously exhibiting the same behavior, this was not considered a separate event. The event would be considered active as long as at least one ant was exhibiting the behavior. We tested all three treatments with each colony.

### Statistical Analysis

#### Lipidomics Statistical Analysis

All lipidomics datasets were first subjected to quality control processing, which included an initial outlier identification step using the algorithm RMD-PAV (47), which specifically performs a robust Principal Component Analysis (PCA) and identifies outliers via distributional properties of the lipid measurements within each sample (Average Pearson correlation within sample type group, median absolute deviation, skew and kurtosis). From the robust PCA a robust Mahalanobis distance is computed and the statistical score associated with each sample is computed from a chi-square distribution with 3 degrees of freedom. Using RMD-PAV we found no outliers. Additionally, the data had very few missing values and thus no lipid identifications were removed either as part of the quality control processing. Quality control processing also evaluated if there was a garden effect, meaning that the samples collected within a garden were more similar than samples collected from different gardens. We computed a standard Pearson correlation values across all pairs of samples as our metric of similarity and compared the within garden correlation values (n1 = 18) with the across garden correlations (n2 = 135) using a two-sample t-test and did not observe a difference based on garden *(p~0.894)*; Pearson correlation of 0.879 and 0.876 for within and across samples, respectively. The requirements of a two-sample t-test were tested for normality with a Kolmogorov-Smirnov test *(p~0.587)* and equal variance with a two-sample F-test *(p~0.125)*. This is confirmed in **Figure 2** with no obvious clustering outside sample levels by garden.

A standard Analysis of Variance (ANOVA) was used to evaluate each lipid for a statistical difference using the factor level (top, middle, and bottom region of the garden) for the first comparison with a Tukey’s post-hoc test to compare the individual levels to one another where each of the three levels contained 6 replicates. These 6 biological replicates provided parallel measurements across 6 distinct fungal garden samples. These were further adjusted with a Benjamini-Hochberg false discovery rate calculation to adjust for the multiple tests being performed across lipids. Adjusted p-values were computed within each negative and positive model lipid dataset and combined for summarization and biological interpretation. **Figure 2** depicts the Tukey’s post-hoc garden top vs garden bottom test results for the 59 significant lipids at a p-value threshold of 0.05. In **Figure 2**, 51 of the 59 lipids were also significant at a Benjamini-Hochberg Tukey’s post-hoc adjusted p-value threshold of 0.05 (Data file S2). The assumptions of the ANOVA model, normality and equal variance, were tested with a Kolmogorov-Smirnov and Bartlett test, respectively. None of the lipids failed to pass the test of normality at either a raw or adjusted p-value. There were only 5 lipids that did not have equal variance at an adjusted p-value, verifying the ANOVA model assumptions for the majority of the lipids and did not warrant using non-parametric statistics for the analysis.

To test for differences between the middle region of the garden and gongylidia for the second comparison (**Figure 3**) a paired t-test was employed, again each factor contained 6 biological replicates. There were 14 (6.2%) lipids that were either absent or nearly absent from one sample type, but not the other (i.e., a lipid was detected in 6 middle garden samples and one or none of the gongylidia samples). To analyze these a G-test was utilized (48) and fold-changes were set to the difference in the counts between the two gardens. Again, the t-test assumptions were evaluated, normality of the difference with a Kolmogorov-Smirnov test. None of the lipids failed to pass the test of normality at either a raw or a Benjamini-Hochberg adjusted p-value. **Figure 3** depicts the significant results for the paired t-test comparing garden middle vs gongylidia for the 116 lipids at a p-value threshold of 0.05. In **Figure 3**, 109 of the 116 lipids were also significant at an adjusted p-value threshold of 0.05 (Data file S3).

#### Statistical data included in **Figure 2** and **Figure 3**

We evaluated if multivariate differences in the lipid profiles between garden components were observable by unsupervised learning, PCA. We conducted one PCA to compare the layers of the garden (**Fig. 2**) and another for the comparison between gongylidia and the middle of the garden (**Fig. 3**). In both cases, we used the function prcomp in the stats package of R with the scale option. To determine if the distance between groups was significantly different than random, we used ANOSIM and PERMANOVA, both in R (49). **Figure 2** contains the log_2_ intensity values for lipids significantly changing (p-value < 0.05 and adjusted p-values < 0.05; exceptions are denoted by #) between the top and bottom regions of the garden where green denotes lipids relatively increased in the top of the garden and orange font denotes lipids relatively increased in the bottom of the garden. **Figure 3** contains the log_2_ intensity values for lipids significantly changing (p-value < 0.05 and adjusted p-values < 0.05; exceptions are denoted by #) between the gongylidia and the middle region of the fungus garden from where they were harvested where teal denotes lipids relatively increased in the garden and purple font denotes lipids relatively increased in the gongylidia. Individual raw p-values and adjusted p-values for each lipid can be found in Data file S2 and Data file S3.

#### Behavior experiment Statistical Analysis

For the behavior experiment, we did not compare the various treatments to each other, but only to the control. Despite being the same age and size, the colonies differed in terms of activity levels and number of ants, so we corrected for this by comparing the response to the treatment disc to the control. For the duration measurements we evaluated the difference in time between treatment and control, and for the number of pickups and aggression events we used the proportion of times the event happened between treatment discs and control discs. For the duration of time we conducted a one sample t-test, where a mean different from zero indicated a significant difference between the treatment and the control, and for the pickup and aggression events we performed a test of proportions, where a proportion different from 0.5 indicated a significant difference (49).

## Acknowledgments

This work was funded by the U.S. Department of Energy (DOE), Office of Science, Office of Biological and Environmental Research (BER), Early Career Research Program Award to KEBJ; Great Lakes Bioenergy Research Center (DOE BER DE-FC02-07ER64494); National Institute of Food and Agriculture, United States Department of Agriculture, under ID number 1003779; and National Institutes of Health (NIH grant U19TW009872). This research used capabilities developed by NIH National Institute of Child Health and Human Development (R21 HD084788), NIH National Institute of Environmental Health Sciences (R01 ES022190) and the Pan-omics program (funded by DOE BER Genome Sciences Program). This work was performed in the W. R. Wiley Environmental Molecular Sciences Laboratory (EMSL) (grid.436923.9), a DOE Office of Science User Facility sponsored by the Office of Biological and Environmental Research and located at Pacific Northwest National Laboratory (PNNL). PNNL is a multiprogram national laboratory operated by Battelle for the Department of Energy (DOE) under Contract DE-AC05-76RL01830. The funders had no role in study design, data collection and interpretation, or the decision to submit the work for publication

## Author contributions

Conceived of the project (KEBJ, ESB, CC, LK), conducted experiments (LK, CN, KEBJ, ESB) performed statistical analysis (BJMWR, LK), analyzed data (LK, KEBJ, ESB, JEK, FBS) designed sampling methodology (LK, KEBJ, CC), provided resources and facilities (KEBJ, ESB, CC, MSL, RDS) and wrote the manuscript (LK, KEBJ, ESB).

## Competing interests

There are no competing interests.

## Data and materials availability

All data are available in the Supplementary Materials. Raw lipidomic datasets are deposited on Mass Spectrometry Interactive Virtual Environment (MassIVE). After the manuscript is published and has a PubMed ID, the data will be made public and available on MassIVE; [https://massive.ucsd.edu] accession number MSV000084520.

## SUPPLEMENTARY MATERIALS LEGENDS

**Figure S1: Heatmaps representing the relative log_2_ expression level for lipids in the leaf food source and the top, middle, and bottom strata of six leaf-cutter ant fungal gardens**. Each column represents the relative abundance of a unique lipid and the heatmaps are scaled by column. Lipids significantly increased in the top are denoted with green font; lipids significantly increasing in the bottom are denoted with orange font (p-values < 0.05 were deemed significant); lipids not significantly changing across the top to the bottom of the garden are denoted with grey font; _A and _B denote structural isomers. Diacylglycerophosphocholines (PC), monoacylglycerophosphocholines (LPC), sulfoquinovosyldiacylglycerol (SQDG), Monogalactosyldiacylglycerol (MGDG), Cardiolipin (CL), phosphatidic acid (PA), Hexosyl-Ceramide (HexCer), Ceramide (Cer), Diacylglycerols (DG), diacylglycerophosphoglycerol (PG), monoacylglycerophosphoglycerol (LPG), diacylglycerophosphoserine (PS), diacylglycerophosphoethanolamines (PE), monoacylglycerophosphoethanolamines (LPE), diacylglycerophosphoinositols (PI). Lipid abbreviations show the total number of acyl chain carbons: total number of double bonds. ‡, 18:2 containing lipids significantly increased in the bottom of the garden; *, 18:3 containing lipids significantly increased in the top of the garden.

**Figure S2: Heatmaps representing the relative log_2_ expression level for lipids in the leaf food source and the gongylidia and surrounding fungal garden of six leaf-cutter ant fungal garden ecosystems**. Each column represents the relative abundance of a unique lipid and the heatmaps are scaled by column. Lipids significantly enriched in the gongylidia compared to the surrounding fungal garden are denoted with purple font; lipids significantly enriched in the fungal garden are denoted with teal font (p-values < 0.05 were deemed significant); lipids not significantly changing between the gongylidia and the surrounding fungal garden are denoted with grey font; _A and _B denote structural isomers. Digalactosyldiacylglycerol (DGDG), sulfoquinovosyldiacylglycerol (SQDG), Monogalactosyldiacylglycerol (MGDG), diacylglycerophosphoglycerol (PG), monoacylglycerophosphoglycerol (LPG), Diacylglycerophosphocholines (PC), monoacylglycerophosphocholines (LPC), mannosylinositol phosphorylceramide (MIPC), Ceramide phosphoinositol (PI-Cer), diacylglycerophosphoinositols (PI), phosphatidic acid (PA), lysophosphatidic acid (LPA), Hexosyl-Ceramide (HexCer), Ceramide (Cer), diacylglycerophosphoethanolamines (PE), monoacylglycerophosphoethanolamines (LPE), Triacylglycerols (TG), Diacylglycerols (DG). Lipid abbreviations show the total number of acyl chain carbons: total number of double bonds. *, 18:3 containing lipids significantly decreased in the gongylidia; ‡, 18:2 containing lipids significantly increased in the gongylidia.

**Figure S3: *Atta* ant behavior in response to alpha-linolenic acid (18:3) in comparison to control**. Labels above panels represent the different colonies that were tested. Tracks represent the times of event occurrences and their durations.

**Figure S4: *Atta* ant behavior in response to linoleic acid (18:2) in comparison to control**. Labels above panels represent the different colonies that were tested. Tracks represent the times of event occurrences and their durations.

**Figure S5: *Atta* ant behavior in response to oleic acid (18:1) in comparison to control**. Labels above panels represent the different colonies that were tested. Tracks represent the times of event occurrences and their durations.

**Data file S1: Proteomic identification of lipid associated proteins**. Quantities of protein spectra related to lipid metabolism identified throughout the fungus garden, from the dataset produced for Aylward et al. 2015.

**Data file S2: Lipid comparisons of top, middle, and bottom fungal garden regions**. Lipidomic intensity data and statistical analysis results for comparisons across the top, middle, and bottom fungal garden regions.

**Data file S3: Lipid changes between fungal garden and gongylidia**. Lipidomic intensity data and statistical analysis results for comparisons across the middle fungal garden and gongylidia.

**Movie S1: Aggressive behaviors.** Discs labeled D are the negative control infused with only ethanol. Discs labeled A are infused with ethanol and alpha-linolenic acid. In the first 34 s, ants can be seen near or on both types of discs with their mandibles open, moving very slowly or standing still. In the second part of the movie, an ant can be seen lunging and biting at the alpha-linolenic acid infused discs. Movie can be viewed at https://youtu.be/fq6SVF-8v6M.

**Movie S2: Interest/attractive behavior**. Discs labeled D are the negative control infused with only ethanol. Discs labeled B are infused with ethanol and linoleic acid. In the first 12 s of the movie, ants can be seen approaching the linoleic acid discs, and waving their antennae. In the second part of the movie, ants are inspecting the linoleic acid discs and one ant picks up a disc at 30 s. Movie can be viewed at https://youtu.be/unUL-ets3Vk.

## Notes

### Competing Interest Statement

The authors have declared no competing interest.

## References

1. Merrill AL, Watt BK. 1973. Energy Value Of Foods - Basis and Derivation.

2. Yeagle PL. 1989. Lipid regulation of cell membrane structure and function. FASEB J 3:1833–42.

3. Waters CM, Bassler BL. 2005. Quorum sensing: Cell-to-cell communication in bacteria. Annual Review of Cell and Developmental Biology 21:319–346.

4. Lebeer S, Vanderleyden J, De Keersmaecker SCJ. 2010. Host interactions of probiotic bacterial surface molecules: comparison with commensals and pathogens. Nature Reviews Microbiology 8:171–184.

5. Selander E, Kubanek J, Hamberg M, Andersson MX, Cervin G, Pavia H. 2015. Predator lipids induce paralytic shellfish toxins in bloom-forming algae. Proceedings of the National Academy of Sciences of the United States of America 112:6395–6400.

6. Kearns DB, Shimkets LJ. 2001. Lipid chemotaxis and signal transduction in Myxococcus xanthus. Trends in Microbiology 9:126–129.

7. Ketter E, Randall G. 2019. Virus Impact on Lipids and Membranes. Annual Review of Virology 6:319–340.

8. Magalhaes DM, Da Silva ITFA, Borges M, Laumann RA, Blassioli-Moraes MC. 2019. Anthonomus grandis aggregation pheromone induces cotton indirect defence and attracts the parasitic wasp Bracon vulgaris. Journal of Experimental Botany 70:1891–1901.

9. Costa AN, Vasconcelos HL, Vieira Neto EHM, Bruna EM. 2008. Do herbivores exert top down effects in Neotropical savannas? Estimates of biomass consumption by leaf cutter ants. Journal of Vegetation Science 19:849–854.

10. Hölldobler B, Wilson EO. 2009. The superorganism: the beauty, elegance, and strangeness of insect societies. WW Norton & Company, New York, NY.

11. Meyer ST, Leal IR, Tabarelli M, Wirth R. 2011. Ecosystem engineering by leaf-cutting ants: nests of *Atta cephalotes* drastically alter forest structure and microclimate. Ecological Entomology 36:14–24.

12. Holldobler B, Wilson EO. 2010. The Leaf-cutter Ants: Civilization by Instinct. W.W. Norton and Company.

13. Aylward FO, Burnum-Johnson KE, Tringe SG, Teiling C, Tremmel DM, Moeller JA, Scott JJ, Barry KW, Piehowski PD, Nicora CD, Malfatti SA, Monroe ME, Purvine SO, Goodwin LA, Smith RD, Weinstock GM, Gerardo NM, Suen G, Lipton MS, Currie CR. 2013. *Leucoagaricus gongylophorus* produces diverse enzymes for the degradation of recalcitrant plant polymers in leaf-cutter ant fungus gardens. Appl Environ Microbiol 79:3770–8.

14. Khadempour L, Burnum-Johnson KE, Baker ES, Nicora CD, Webb-Robertson BM, White RA, 3rd, Monroe ME, Huang EL, Smith RD, Currie CR. 2016. The fungal cultivar of leaf-cutter ants produces specific enzymes in response to different plant substrates. Mol Ecol 25:5795–5805.

15. Suen G, Teiling C, Li L, Holt C, Abouheif E, Bornberg-Bauer E, Bouffard P, Caldera EJ, Cash E, Cavanaugh A, Denas O, Elhaik E, Fave MJ, Gadau J, Gibson JD, Graur D, Grubbs KJ, Hagen DE, Harkins TT, Helmkampf M, Hu H, Johnson BR, Kim J, Marsh SE, Moeller JA, Munoz-Torres MC, Murphy MC, Naughton MC, Nigam S, Overson R, Rajakumar R, Reese JT, Scott JJ, Smith CR, Tao S, Tsutsui ND, Viljakainen L, Wissler L, Yandell MD, Zimmer F, Taylor J, Slater SC, Clifton SW, Warren WC, Elsik CG, Smith CD, Weinstock GM, Gerardo NM, Currie CR. 2011. The genome sequence of the leaf-cutter ant Atta cephalotes reveals insights into its obligate symbiotic lifestyle. PLoS Genet 7:e1002007.

16. Nagamoto NS, Garcia MG, Forti LC, Verza SS, Noronha NC, Rodella RA. 2011. Microscopic evidence supports the hypothesis of high cellulose degradation capacity by the symbiotic fungus of leaf-cutting ants. Journal of Biological Research-Thessaloniki 16:308–312.

17. Suen G, Scott JJ, Aylward FO, Currie CR. 2011. The Microbiome of Leaf-Cutter Ant Fungus Gardens, p 367–379. *In* de Bruijn FJ (ed), Handbook of Molecular Microbial Ecology II doi:10.1002/9781118010549.ch36. John Wiley & Sons, Inc., Hoboken, NJ, USA.

18. Moreira-Soto RD, Sanchez E, Currie CR, Pinto-Tomás AA. 2017. Ultrastructural and microbial analyses of cellulose degradation in leaf-cutter ant colonies. Microbiology 163:1578–1589.

19. De Fine Licht HH, Schiott M, Rogowska-Wrzesinska A, Nygaard S, Roepstorff P, Boomsma JJ. 2013. Laccase detoxification mediates the nutritional alliance between leaf-cutting ants and fungus-garden symbionts. Proc Natl Acad Sci U S A 110:583–7.

20. Aylward FO, Khadempour L, Tremmel DM, McDonald BR, Nicora CD, Wu S, Moore RJ, Orton DJ, Monroe ME, Piehowski PD, Purvine SO, Smith RD, Lipton MS, Burnum-Johnson KE, Currie CR. 2015. Enrichment and Broad Representation of Plant Biomass-Degrading Enzymes in the Specialized Hyphal Swellings of Leucoagaricus gongylophorus, the Fungal Symbiont of Leaf-Cutter Ants. PLoS One 10:e0134752.

21. Farji-Brener AG, Medina CA. 2000. The Importance of Where to Dump the Refuse: Seed Banks and Fine Roots in Nests of the Leaf-Cutting Ants Atta cephalotes and A. colombica1. Biotropica 32:120–126.

22. Bass M, Cherrett JM. 1995. Fungal Hyphae as a Source of Nutrients for the Leaf-Cutting Ant Atta Sexdens. Physiological Entomology 20:1–6.

23. Shik JZ, Rytter W, Arnan X, Michelsen A. 2018. Disentangling nutritional pathways linking leaf-cutter ants and their co-evolved fungal symbionts using stable isotopes. Ecology 99:1999–2009.

24. Aylward FO, Burnum KE, Scott JJ, Suen G, Tringe SG, Adams SM, Barry KW, Nicora CD, Piehowski PD, Purvine SO, Starrett GJ, Goodwin LA, Smith RD, Lipton MS, Currie CR. 2012. Metagenomic and metaproteomic insights into bacterial communities in leaf-cutter ant fungus gardens. Isme Journal 6:1688–1701.

25. Kyle JE, Zhang X, Weitz KK, Monroe ME, Ibrahim YM, Moore RJ, Cha J, Sun X, Lovelace ES, Wagoner J, Polyak SJ, Metz TO, Dey SK, Smith RD, Burnum-Johnson KE, Baker ES. 2016. Uncovering biologically significant lipid isomers with liquid chromatography, ion mobility spectrometry and mass spectrometry. Analyst 141:1649–59.

26. Schultz TR, Brady SG. 2008. Major evolutionary transitions in ant agriculture. Proc Natl Acad Sci U S A 105:5435–40.

27. Nygaard S, Hu H, Li C, Schiøtt M, Chen Z, Yang Z, Xie Q, Ma C, Deng Y, Dikow RB, Rabeling C, Nash DR, Wcislo WT, Brady SG, Schultz TR, Zhang G, Boomsma JJ. 2016. Reciprocal genomic evolution in the ant-fungus agricultural symbiosis. Nature Communications 7:12233.

28. Kobayashi K. 2016. Role of membrane glycerolipids in photosynthesis, thylakoid biogenesis and chloroplast development. J Plant Res 129:565–580.

29. Conconi A, Miquel M, Browse JA, Ryan CA. 1996. Intracellular Levels of Free Linolenic and Linoleic Acids Increase in Tomato Leaves in Response to Wounding. Plant Physiol 111:797–803.

30. Gordon DM. 1983. Dependence of necrophoric response to oleic acid on social context in the ant, Pogonomyrmex badius. J Chem Ecol 9:105–11.

31. Wilson EO, Durlach NI, Roth LM. 1958. Chemical Releaser of Necrophoric Behavior in Ants. Psyche: A Journal of Entomology 65:108–114.

32. Lopez-Riquelme GO, Malo EA, Cruz-Lopez L, Fanjul-Moles ML. 2006. Antennal olfactory sensitivity in response to task-related odours of three castes of the ant Atta mexicana (Hymenoptera: Formicidae). Physiological Entomology 31:353–360.

33. Buehlmann C, Graham P, Hansson BS, Knaden M. 2014. Desert ants locate food by combining high sensitivity to food odors with extensive crosswind runs. Curr Biol 24:960–4.

34. De Fine Licht HH, Boomsma JJ, Tunlid A. 2014. Symbiotic adaptations in the fungal cultivar of leaf-cutting ants. Nat Commun 5:5675.

35. Kecskeméti S, Szelényi MO, Erdei AL, Geösel A, Fail J, Molnár BP. 2020. Fungal Volatiles as Olfactory Cues for Female Fungus Gnat, Lycoriella ingenua in the Avoidance of Mycelia Colonized Compost. Journal of Chemical Ecology 46:917–926.

36. Holighaus G, Rohlfs M. 2016. Fungal allelochemicals in insect pest management. Applied Microbiology and Biotechnology 100:5681–5689.

37. North RD, Jackson CW, Howse PE. 1999. Communication between the fungus garden and workers of the leaf-cutting ant, Atta sexdens rubropilosa, regarding choice of substrate for the fungus. Physiological Entomology 24:127–133.

38. Sousa KKA, Catalani GC, Gianeti TMR, Camargo RS, Caldato N, Ramos VM, Forti LC. 2020. A Volatile Semiochemical Released by the Fungus Garden of Leaf-Cutting Ants. Florida Entomologist 103:1–8, 8.

39. Green PWC, Kooij PW. 2018. The role of chemical signalling in maintenance of the fungus garden by leaf-cutting ants. Chemoecology 28:101–107.

40. Folch J, Lees M, Sloane Stanley GH. 1957. A simple method for the isolation and purification of total lipides from animal tissues. J Biol Chem 226:497–509.

41. Liebisch G, Fahy E, Aoki J, Dennis EA, Durand T, Ejsing C, Fedorova M, Feussner I, Griffiths WJ, Koefeler H, Merrill AH, Jr., Murphy RC, O’Donnell VB, Oskolkova OV, Subramaniam S, Wakelam M, Spener F. 2020. Update on LIPID MAPS Classification, Nomenclature and Shorthand Notation for MS-derived Lipid Structures. J Lipid Res doi:10.1194/jlr.S120001025.

42. Crowell KL, Slysz GW, Baker ES, LaMarche BL, Monroe ME, Ibrahim YM, Payne SH, Anderson GA, Smith RD. 2013. LC-IMS-MS Feature Finder: detecting multidimensional liquid chromatography, ion mobility and mass spectrometry features in complex datasets. Bioinformatics 29:2804–5.

43. Kyle JE, Crowell KL, Casey CP, Fujimoto GM, Kim S, Dautel SE, Smith RD, Payne SH, Metz TO. 2017. LIQUID: an-open source software for identifying lipids in LC-MS/MS-based lipidomics data. Bioinformatics 33:1744–1746.

44. Pluskal T, Castillo S, Villar-Briones A, Oresic M. 2010. MZmine 2: modular framework for processing, visualizing, and analyzing mass spectrometry-based molecular profile data. BMC Bioinformatics 11:395.

45. Friard O, Gamba M. 2016. BORIS: a free, versatile open-source event-logging software for video/audio coding and live observations. Methods in Ecology and Evolution 7:1325–1330.

46. Hernandez JV, Lopez H, Jaffe K. 2002. Nestmate recognition signals of the leaf-cutting ant Atta laevigata. Journal of Insect Physiology 48:287–295.

47. Matzke MM, Waters KM, Metz TO, Jacobs JM, Sims AC, Baric RS, Pounds JG, Webb-Robertson BJ. 2011. Improved quality control processing of peptide-centric LC-MS proteomics data. Bioinformatics 27:2866–72.

48. Webb-Robertson BJ, McCue LA, Waters KM, Matzke MM, Jacobs JM, Metz TO, Varnum SM, Pounds JG. 2010. Combined statistical analyses of peptide intensities and peptide occurrences improves identification of significant peptides from MS-based proteomics data. J Proteome Res 9:5748–56.

49. Team RC. R: A language and environment for statistical computing. R Foundation for Statistical Computing. Available at: http://www.R-project.org/ (Accessed: 23rd December 2013).

