## Supplementary figures and images for "From plants to ants: Fungal modification of leaf lipids for nutrition and communication in the leaf-cutter ant fungal garden ecosystem"

### Figure S1

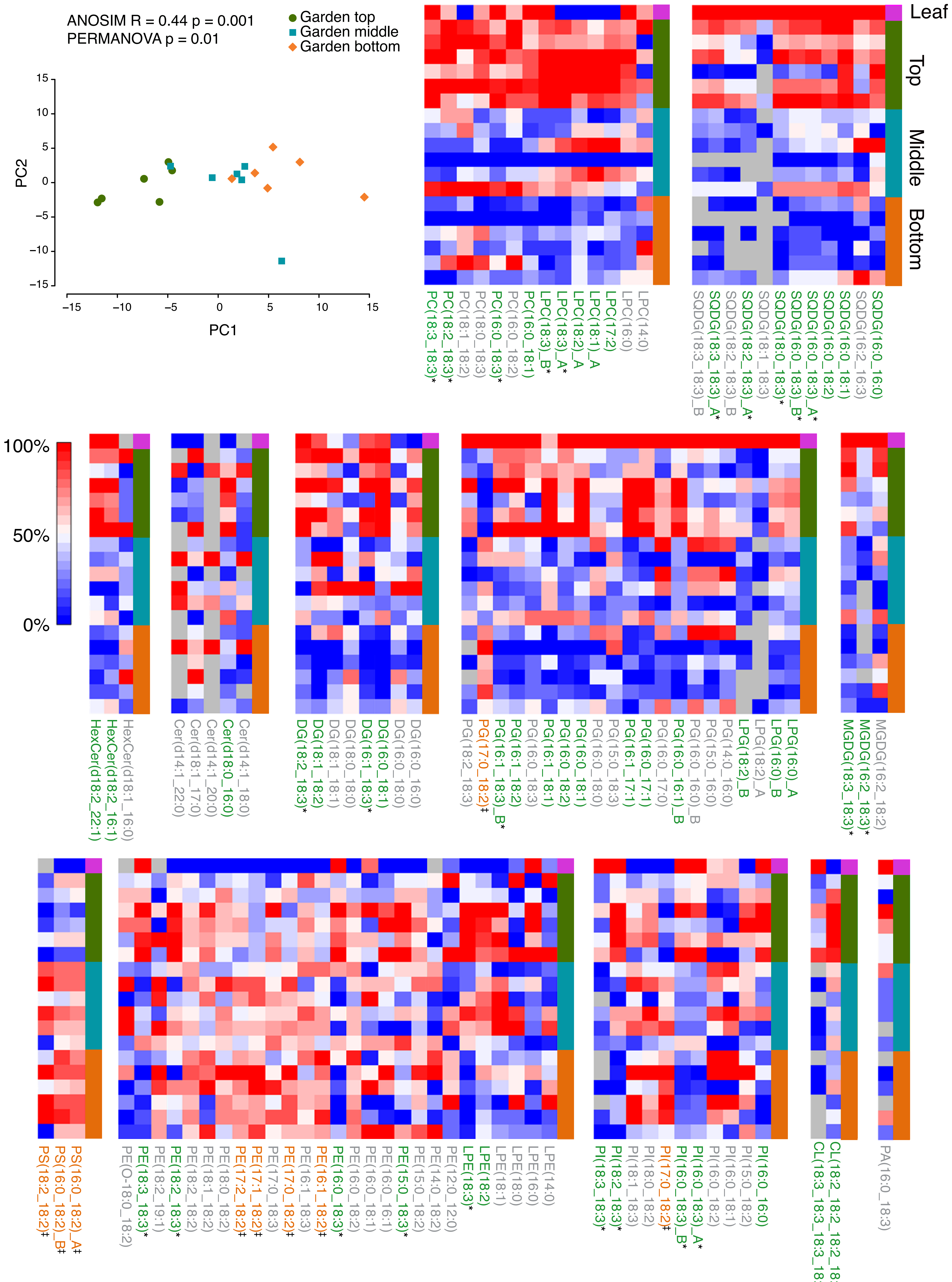

### Figure S2

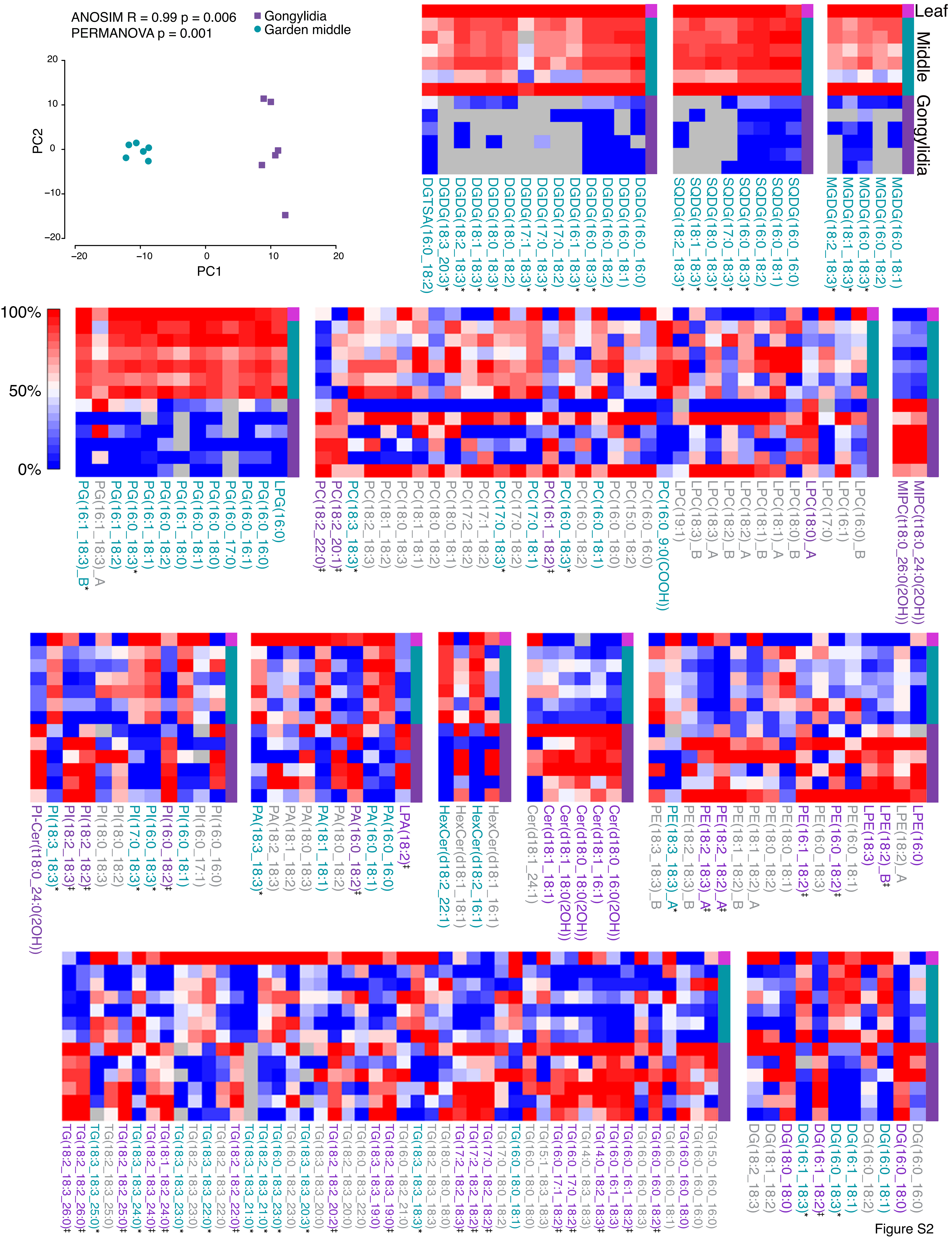

### Figure S3

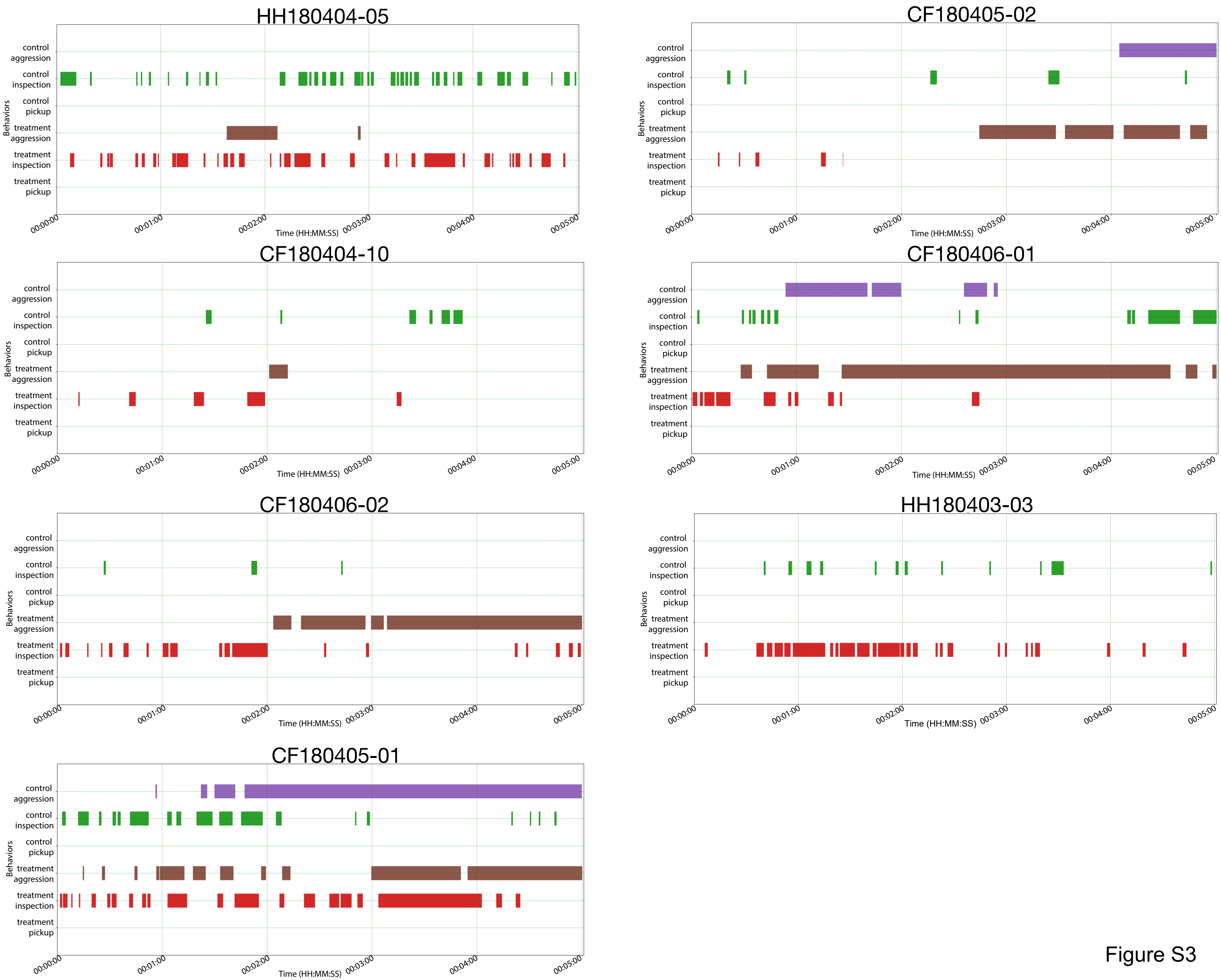

### Figure S4

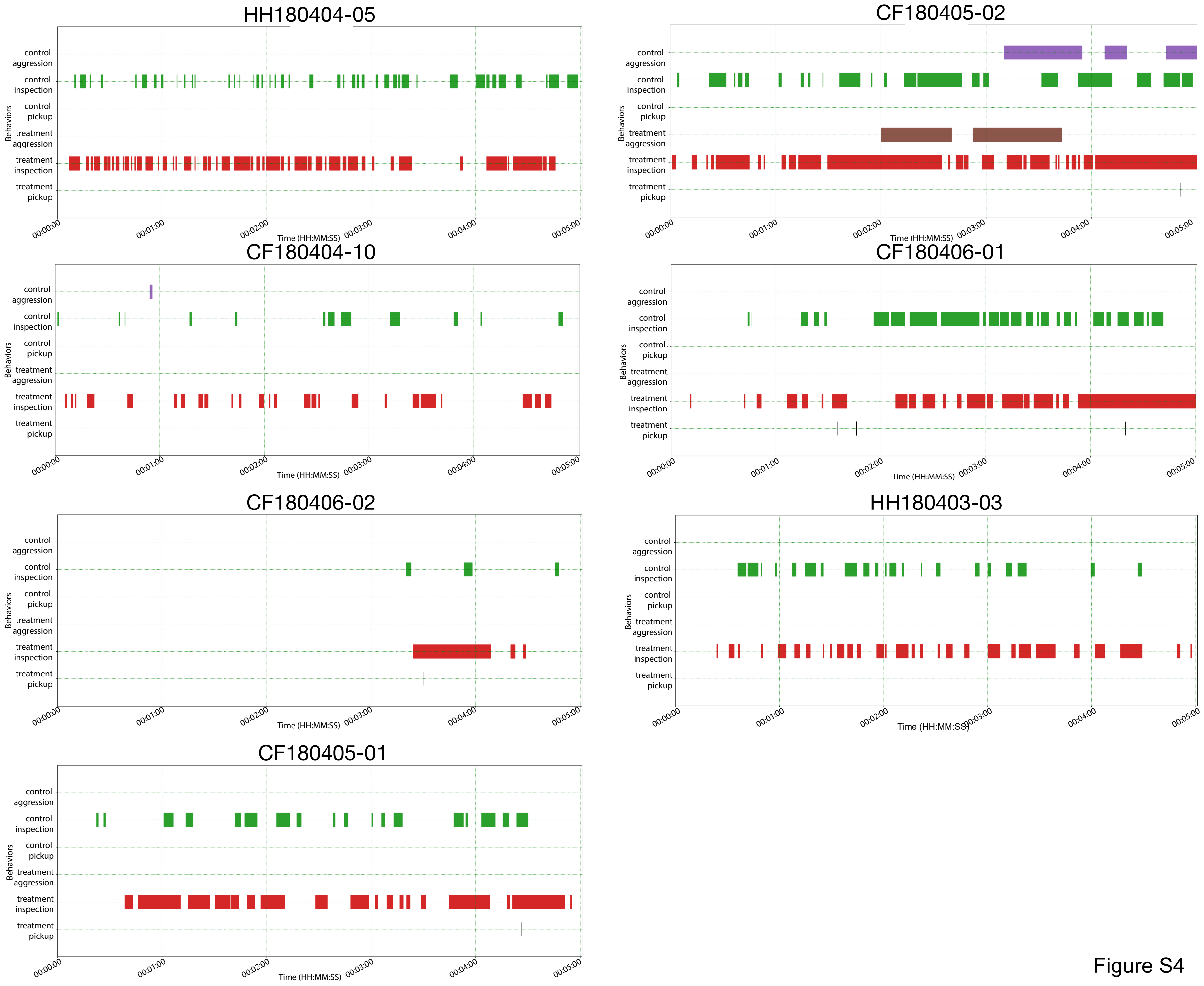

### Figure S5

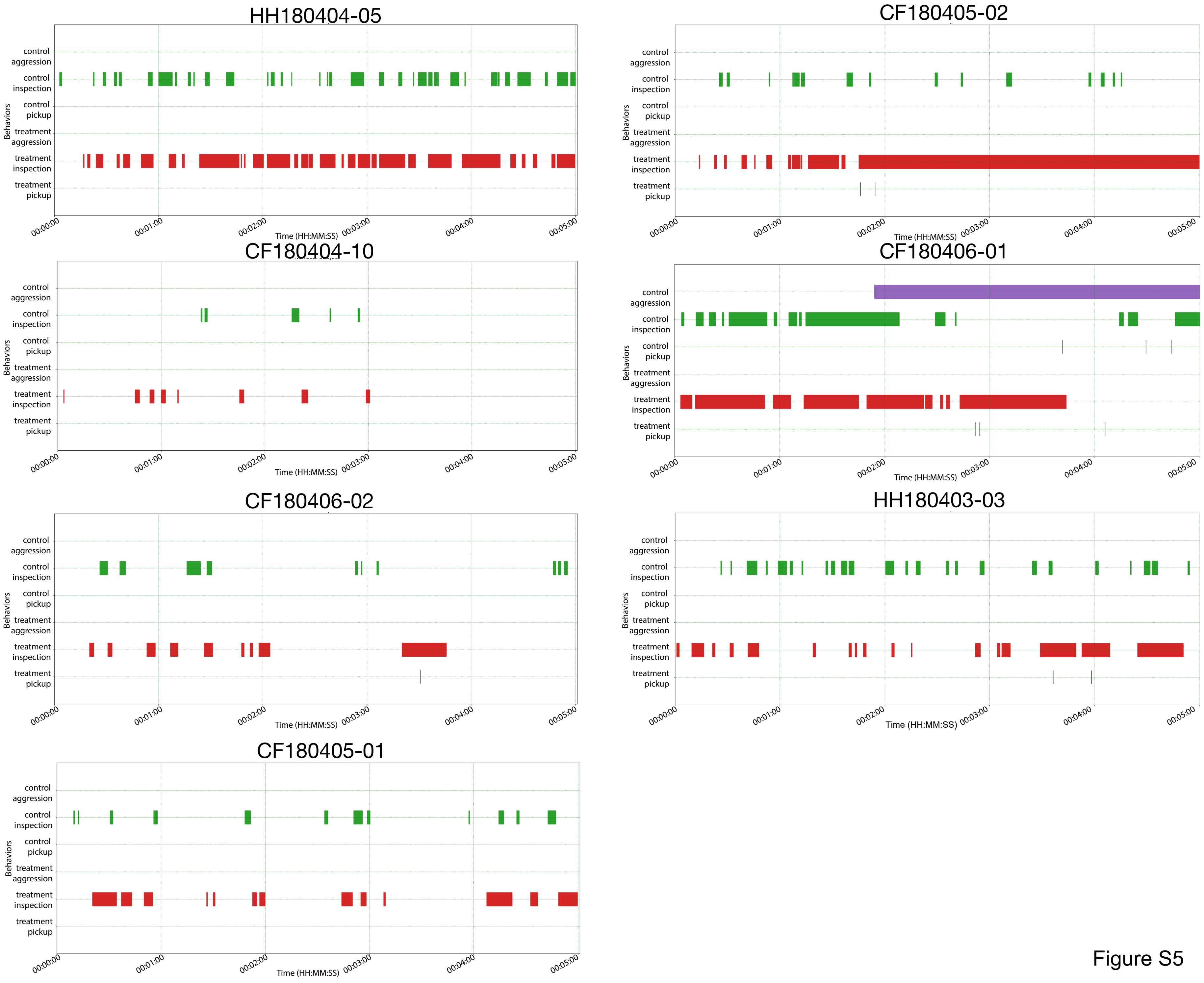
